# Macrophage metabolism in the intestine is compartment-specific and regulated by the microbiota

**DOI:** 10.1101/2021.06.24.449804

**Authors:** Nicholas A. Scott, Melissa A. E. Lawson, Ryan James Hodgetts, Lindsay J. Hall, Elizabeth R. Mann

**Affiliations:** Lydia Becker Institute of Immunology and Inflammation, Division of Infection, Immunity and Respiratory Medicine, School of Biological Sciences, Faculty of Biology, Medicine and Health, University of Manchester, Manchester Academic Health Science Centre, Grafton Street, Manchester, UK; Quadram Institute Bioscience, Norwich Research Park, Norwich, UK; Department of Immunology, Weizmann Institute of Science, Rehovot, Israel

## Abstract

Intestinal macrophages play a vital role in the maintenance of gut homeostasis through signals derived from the microbiota. We previously demonstrated that microbial-derived metabolites can shape the metabolic functions of macrophages. Here, we show that antibiotic-induced disruption of the intestinal microbiota dramatically alters both the local metabolite environment, and the metabolic functions of macrophages in the colon. Broad-spectrum antibiotic administration in mice increased expression of the large neutral amino acid transporter and accordingly, amino acid uptake. Subsequently, antibiotic administration enhanced the metabolic functions of colonic macrophages, increasing phosphorylation of components of mammalian/mechanistic target of rapamycin (mTOR) signalling pathways, increasing expression of genes involved in glycolysis and oxidative phosphorylation (OXPHOS), increasing mitochondrial function and increased levels of ECAR and OCR as a direct measure of glycolysis and OXPHOS. Small bowel macrophages were less metabolically active than in the colon, with macrophage metabolism being independent of the microbiota. Finally, we reveal tissue resident Tim4^+^ CD4^+^ macrophages exhibit enhanced fatty acid uptake alongside reduced fatty acid synthesis compared to their recruited counterparts. Thus the microbiota shapes gut macrophage metabolism in a compartment-specific manner, with important implications for functions when monocyte recruitment and macrophage differentiation.

## INTRODUCTION

Intestinal macrophages in the lamina propria are specialised to remain hyporesponsive to the gut microbiota, while remaining poised to respond to pathogens when required. Under steady state conditions, intestinal macrophages do not respond to bacterial stimulation, such as lipopolysaccharide (LPS)^1,2^. This is thought to be one of the major mechanisms by which harmful immune responses against the intestinal microbiota are avoided. Most macrophages within the intestine are seeded from Ly6C^+^ MHC class II^−^ blood circulating monocytes, which differentiate to an intermediate stage (Ly6C^+^ MHC class II^+^), before maturing to macrophages (Ly6C^−^ MHC class II^+^)^3^. However, Tim4 and CD4 expression identifies macrophages with low turnover from the circulation^4^, although the proportions of these macrophages differ between the small and large intestine and functional differences between recently recruited and long-lived macrophages in the intestine are unclear.

Metabolic pathways provide energy to cells but are also capable of regulating immune function. Immune cell metabolism regulates their function and is dependent on the local availability of nutrients^5–8^. Classically activated macrophages are typically induced by LPS or IFNγ^9^, defined by production of high levels of proinflammatory cytokines in response to pathogens rely on glucose utilization for energy (anaerobic glycolysis). In contrast, alternatively activated macrophages provide growth factors for wound healing and are generated by type 2 cytokines such as IL-4 or IL-13, using oxidative phosphorylation (OXPHOS) generated within mitochondria from the tricarboxylic acid cycle (TCA) as their primary energy source^10,11^. Primary tissue macrophages exhibit properties of both classical and alternative activation, and it is now thought that a spectrum of macrophage phenotypes exist which are shaped by their local tissue niche.

Amino acids, as well as providing the building blocks of proteins and nucleotides, govern cellular metabolism by forming precursors of key regulatory metabolites^12^. Amino acid uptake has been proposed to be important for macrophage function as LPS stimulation of bone marrow-derived macrophages increases expression of Slc7a5, a component of the amino acid transporter LAT1. Subsequent leucine importation through LAT1 regulates proinflammatory cytokine production by macrophages^13^. This is of particular relevance to the intestine due to high levels of amino acids from intake of dietary protein. It was recently demonstrated that deletion of Slc3a2, a subunit of amino acid transporter LAT1 in CX3CR1^+^ gut macrophages reduces their numbers and impedes monocyte differentiation to mature MHC class II^+^ macrophages^14^.

The gut microbiota generates metabolites that regulate the intestinal immune system, with arguably the most extensively studied being short-chain fatty acids^15–18^. We and others have demonstrated the SCFA butyrate induces metabolic rewiring in intestinal macrophages towards oxidative phosphorylation^19^ and away from glycolysis^20^. Nonetheless, the overall contribution of the microbiota to macrophage metabolism in the intestine remains unclear. Here, we demonstrate that antibiotic-induced disruption of the intestinal microbiota enhances macrophage amino acid uptake and metabolic capacity, with upregulation of glycolysis as well as OXPHOS pathways. These changes were not SCFA-dependent and limited to colonic macrophages only. Indeed, the metabolic capacity of macrophages in the small intestine was diminished compared to that of colonic macrophages. The importance of considering macrophage subsets in the intestine is demonstrated with fatty acid uptake being a defining characteristic of long-lived Tim4^+^CD4^+^ colonic tissue resident macrophages.

## RESULTS

### Antibiotic-treatment increases amino acid uptake and expression of the LAT1 transporter on colonic macrophages

Given the high levels of amino acids in the intestine and contribution of components of the amino acid transporter LAT1 to the differentiation of intestinal macrophages^14^, we first examined amino acid transport in monocytes and macrophages of the intestine. Leucine uptake promotes pro-inflammatory cytokine production in macrophages^13^, with importation of leucine into cells occurring via the transporter LAT1, a heterodimer of CD98 and Slc7a5. We assessed the expression of CD98 and Slc7a5 in the CD11b^+^CD64^+^ monocyte and macrophage subsets from the colon. Protein and transcript levels of CD98 and Slc7a5 respectively, increased significantly as monocytes (Ly6C^+^MHCcII^−^) matured to intermediate monocytes (Ly6C^+^MHCcII^+^) and increased further for CD98 following macrophage differentiation (Ly6C^−^MHCcII^+^) (Fig. 1A and 1B), inferring amino acid transport may be enhanced during maturation of monocytes.

**Figure 1:**
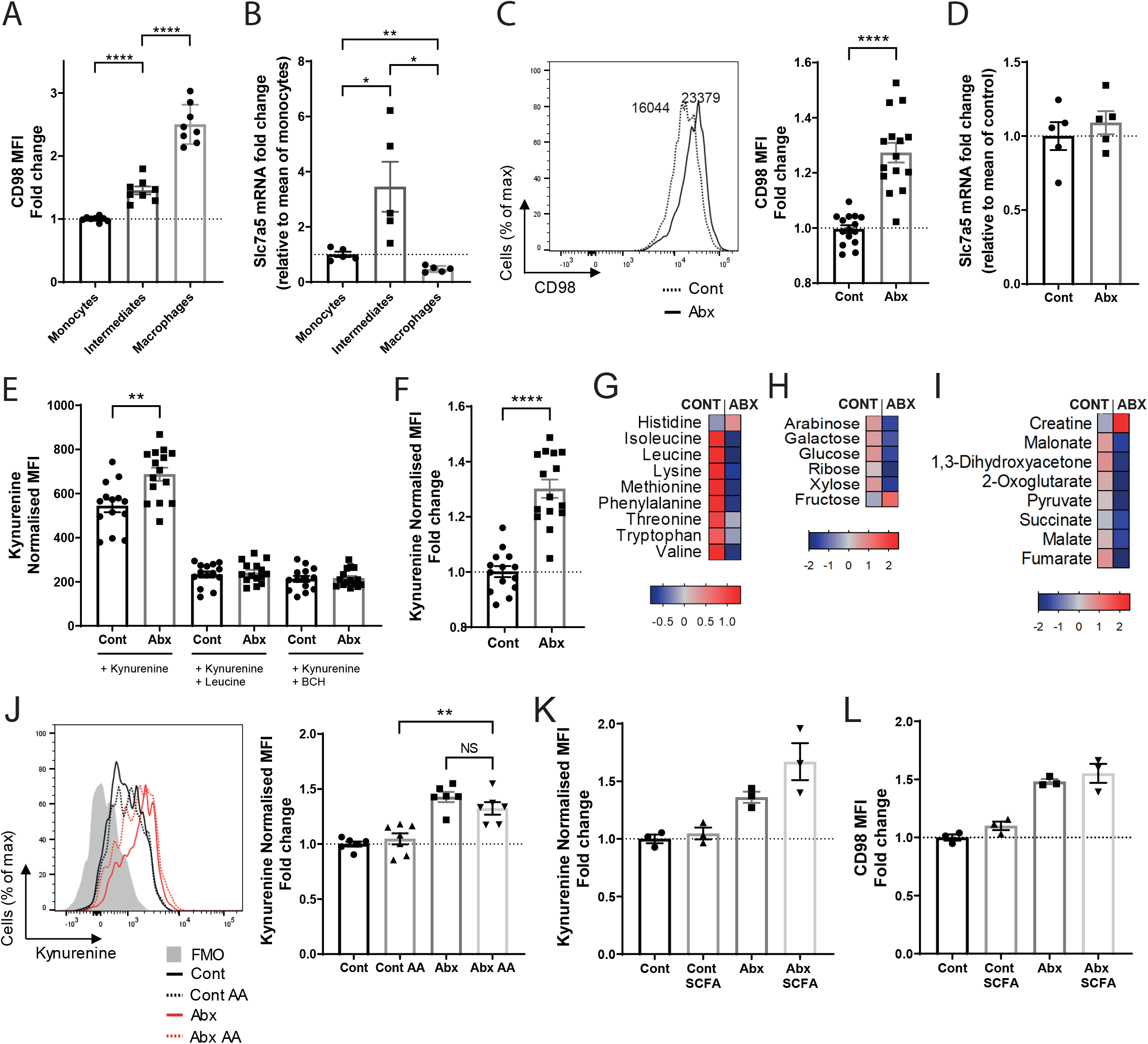
Antibiotic treatment alters amino acid transport in large intestinal macrophages. **(A-F)** Mice were treated with antibiotic or control water and colonic monocyte/macrophage populations (gated from Live CD45^+^Ly6G^−^CD11b^+^CD64^+^) characterised by flow cytometry or sorted for qPCR. **(A)** Mean fluorescent intensity of CD98 protein expression at steady state on monocytes (Ly6C^+^MHC cII^−^), intermediates (Ly6C^+^MHC cII^+^) and macrophages (Ly6C^−^MHC cII^+^). Fold change is relative to mean monocyte readings; n=8. **(B)** Slc7a5 mRNA expression on same subsets shown in **A**. Fold change is relative to mean monocyte readings; n=5. **(C)** Mean fluorescent intensity of CD98 protein expression on macrophages (Ly6C^−^MHC cII^+^). Fold change is relative to mean control readings; n=15. **(D)** Slc7a5 mRNA expression on Ly6C^−^MHC cII^+^ macrophages. Fold change is relative to mean of control group; n=5. (**E**) Intestinal cell suspensions were treated with 200μM kynurenine and cell uptake measured on Ly6C^−^MHC cII^+^ intestinal macrophages at 450nm in the presence or absence of competitive leucine (5mM) or the System L transport inhibitor BCH (10mM). MFI readings were normalised to no kynurenine samples; n=15. **(F)** Data from (E) (kynurenine alone) expressed as fold change relative to control treated, n=15. **(G-I)** Mice were treated with antibiotic or control water and faecal pellets analysed by nuclear magnetic resonance (NMR) spectroscopy. Pooled data depicting heatmaps made from Z-scores (relative). **(G)** Fold change in selected sugars. **(H)** Fold change in selected amino acids. **(I)** Fold change in selected energy metabolites. **(J-L)** In addition to control and antibiotic water alone, separate cohorts of mice were treated with control and antibiotic water supplemented with amino acids (AA) (leucine, lysine and glutamine) **(J)** (n=6-7), or short chain fatty acids (SCFA) comprising butyrate (40mM), acetate (67.5mM) and propionate (25mM) **(K)** (n=3). MFI readings are normalised to no kynurenine samples and fold change expressed as mean of control samples. **(L)** Mean fluorescent intensity of CD98 protein expression on colonic macrophages from **K**. Fold change is normalised to mean control readings; n=3. *p<0.05, **p<0.01, ****p<0.0001, NS: non-significant, Student’s t-test.

To assess whether components of amino acid transport pathways may be modulated by the microbiota, we disrupted the gut microbiota by treating mice with a broad-spectrum antibiotic cocktail for seven days. CD98 was upregulated on macrophages from antibiotic-treated mice compared to controls (Fig. 1C), but Slc7a5 expression was not significantly changed (Fig. 1D). Uptake of amino acids was quantified by measuring importation of kynurenine, a florescent compound selectively transported by LAT1^21^, assessed by flow cytometry in CD11b^+^CD64^+^MHC cII^+^Ly6C^−^ colonic macrophages. Macrophages from antibiotic-treated mice demonstrated increased kynurenine (amino acid) uptake (Fig. 1E and 1F) which the addition of competing leucine or the System L inhibitor, 2-amino-2-norbornanecarboxylic acid (BCH), blocked (Fig.1E).

To assess whether enhanced uptake of amino acids following antibiotic treatment may be a result of changes in local levels of amino acids, we assessed the local levels of amino acids and energy related products by analysing faecal pellets using nuclear magnetic resonance spectroscopy. Antibiotic treatment depleted levels of amino acids in the colon (Fig. 1G) including leucine which is important for the production of proinflammatory cytokines by monocytes and macrophages^13^. In addition, glucose, galactose and ribose were all reduced, among other sugars (Fig. 1H), and several other key metabolites declined, such as succinate, generated during mitochondrial respiration in the tricarboxylic acid cycle (TCA), and pyruvate, generated by glycolysis. Interestingly, one of the few molecules elevated is creatine, a catabolite of arginine recently shown to promote M(IL-4) polarisation^22^. Collectively these data demonstrate that disruption of the microbiota in the gut drastically changes the availability of local metabolites that have been shown to modulate macrophage function.

To investigate whether reduced availability of amino acids following antibiotics may account for enhanced amino acid uptake mice were administered antibiotic or control water supplemented with amino acids (leucine, glutamine and lysine). Within both the control and antibiotic groups, the addition of amino acids had no effect on kynurenine (amino acid) uptake in macrophages (Fig. 1J), suggesting local concentrations of amino acid in vivo does not alter macrophage transport. The concentrations of short chain fatty acids (SCFA) including butyrate, propionate and acetate are also decreased with the same antibiotic regimen, with SCFAs having potent effects on macrophage function^19^. To investigate whether changes in local SCFA levels regulate amino acid transport, SCFAs were administered to antibiotic-treated and control mice to examine their effect on amino acid uptake and CD98 expression. Addition of SCFA did not change kynurenine transport (Fig. 1K) or CD98 protein levels (Fig. 2L) in colonic macrophages. These data demonstrate antibiotic-induced disruption of the microbiota alters amino acid uptake in colonic macrophages by a mechanism indirect of local availability of amino acids and SCFAs.

**Figure 2:**
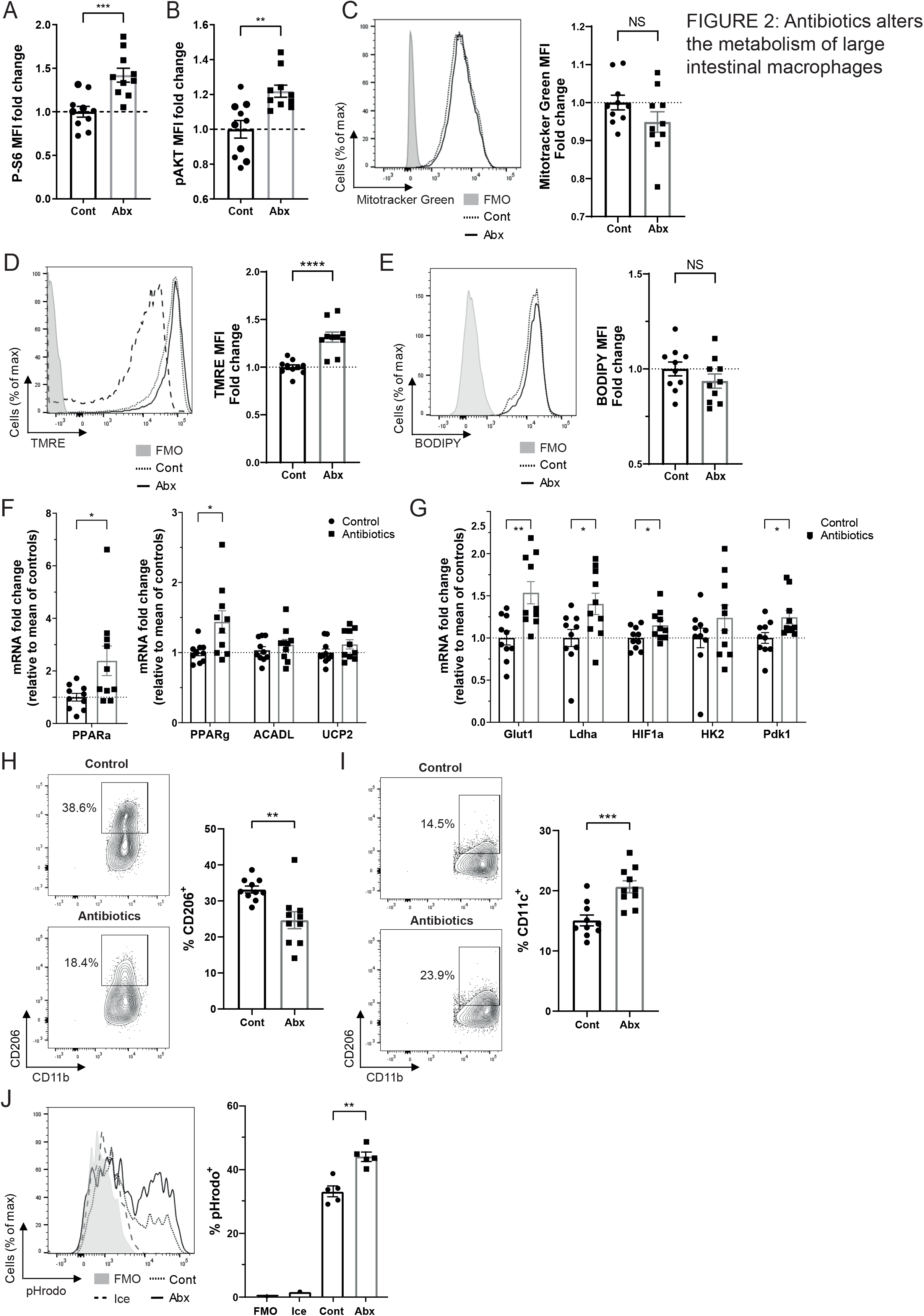
Antibiotics alters the metabolism of macrophages within the large intestine. Mice were treated with antibiotic or control water and large intestinal cell suspensions isolated. **(A-E)** Macrophages (LiveCD45^+^SiglecF^−^Ly6G^−^CD11b^+^CD64^+^Ly6C^−^MHC cII^+^) were identified by flow cytometry for gating. **(A-B)** Phosphoflow for phosphorylated Serine-6-Kinase **(A)** and phosphorylated AKT **(B)**. Fold change is normalised to control samples; n=10. **(C)** Cells were treated with 50nM Mitotracker Green and staining measured at 530nM. n=10. **(D)** Cells were treated with 25nM of TMRE and fluorescence measured at 586nM. n=10. **(E)** Cells were treated with 5μM BODIPY FL C16 in fatty acid free media, and uptake measured at 530nM, n=10. **(F-G)** Large intestinal macrophages (LiveCD45^+^SiglecF^−^Ly6G^−^ CD11b^+^CD64^+^) were sorted by flow cytometry and mRNA expression of glycolytic-related (**F**) and oxidative phosphorylation-related **(G)** genes were quantified by real-time PCR. Fold change is relative to mean of control samples; n=10. **(H-I)** CD206 **(H)** and CD11c **(I)** expression and on Ly6C^−^MHCcII^+^ macrophages by flow cytometry; n-10. **(J)** Cell suspensions were analysed with the pHrodo phagocytosis assay and uptake on macrophages measured at 561nM; n=5.*p<0.05, **p<0.01, ***p<0.001, ****p<0.0001, NS: non-significant, Student’s t-test.

### Antibiotics enhances the metabolic capacity of colonic macrophages

Modulation of cellular metabolism is mediated through mTOR (mechanistic/mammalian target of rapamycin), with amino acids (in particular leucine) enhancing the activity of mTOR complex 1 (mTORC1) through the acetylation of raptor by its metabolite acetyl-coenzyme A^23^. This complex functions as a key nutrient sensing pathway that permits macrophages to regulate cell growth and metabolism. To investigate whether increased amino acid uptake in intestinal macrophages results in increased mTORC1 activity, we analysed phosphorylation of S6-kinase (S6), which is phosphorylated downstream of mTORC1. Colonic macrophages isolated from antibiotic-treated mice had higher levels of phosporylated-S6 than macrophages from control animals (Fig. 2A). mTORC1 is also activated by growth factors and cytokine signalling, through the PIP3K-Akt (class I phosphoinositide 3-kinase-Akt) pathway upstream of mTORC1. Accordingly, we also revealed increased phosphorylation of Akt in gut macrophages from antibiotic-treated mice (Fig. 2B).

We then measured features of glycolysis and oxidative phosphorylation (OXPHOS) following antibiotic treatment, using a range of experimental cellular assays. Although there was no difference in mitochondrial mass and fatty acid uptake (Fig. 2C & 2E), antibiotics significantly enhanced the mitochondrial function of colonic macrophages (Fig. 2D), suggestive of increased levels of OXPHOS. Gene expression of OXPHOS regulatory genes were evaluated by qPCR in CD11b^+^ CD64^+^ colonic macrophages. Macrophages from antibiotic-treated mice had elevated expression of two enzymes involved in transcriptional regulation of fatty acid oxidation (*Pparα* and *Pparγ*) (Fig. 2F), and heightened expression of key glycolysis genes, including *Glut1* (primary glucose transporter), *Ldha* (lactate dehydrogenase, catalyses the final step of glucolysis), *Hif-1a* (hypoxia-inducible factor 1-alpha, a key transcription factor that targets glucose uptake and glycolysis) and *Pdk1* (pyruvate dehydrogenase kinase 1, inhibits TCA cycle; Fig. 2G). Despite OXPHOS being a characteristic of alternatively activated macrophages^24^ the mannose receptor (CD206), a marker of alternative activation, was downregulated in macrophages from antibiotic-treated mice (Fig. 2H) and CD11c, a marker of classical activation, was significantly increased (Fig. 2I). Corresponding with enhanced characteristics of classical activation which is associated with glycolysis^25–27^ antibiotic-induced microbiota-disruption increased the phagocytic activity of intestinal macrophages (Fig. 2J).

To confirm whether antibiotic induced microbial disruption shaped glycolysis and OXPHOS directly, we assessed the metabolic functions of sorted CD11b^+^ CD64^+^ colonic macrophages using Seahorse^TM^ technology. Pooled cells from antibiotic-treated mice exhibited increased extracellular acidification rates (ECAR) as a measure of glycoysis (Fig. 3A) across independent experiments (Fig. 3B), as well as increased oxygen consumption rates (OCR) (Fig. 3C) and maximal mitochondrial respiration as a measure of OXPHOS^28^ (Fig. 3D). The addition of fatty acid oxidation inhibitor etomoxir during the mitochondrial stress test allows for the assessment of the dependence of cells on fatty acid oxidation/lipid metabolism during cellular respiration. The fold change decrease in OCR induced by etomoxir was higher in macrophages isolated from antibiotic-treated mice, indicating a higher dependency of these cells on fatty acid oxidation than their control counterparts (Fig. 3E), as opposed to lipid uptake (Fig. 2E). These data suggest colonic macrophages become increasingly metabolically active in response to antibiotics, demonstrating increased levels of glycolysis, OXPHOS and a higher dependency for fatty acid oxidation. In terms of associating metabolic function with activation, these data are in keeping with our previous studies demonstrating antibiotic treatment induces pro-inflammatory changes in intestinal macrophages^19^.

**Figure 3:**
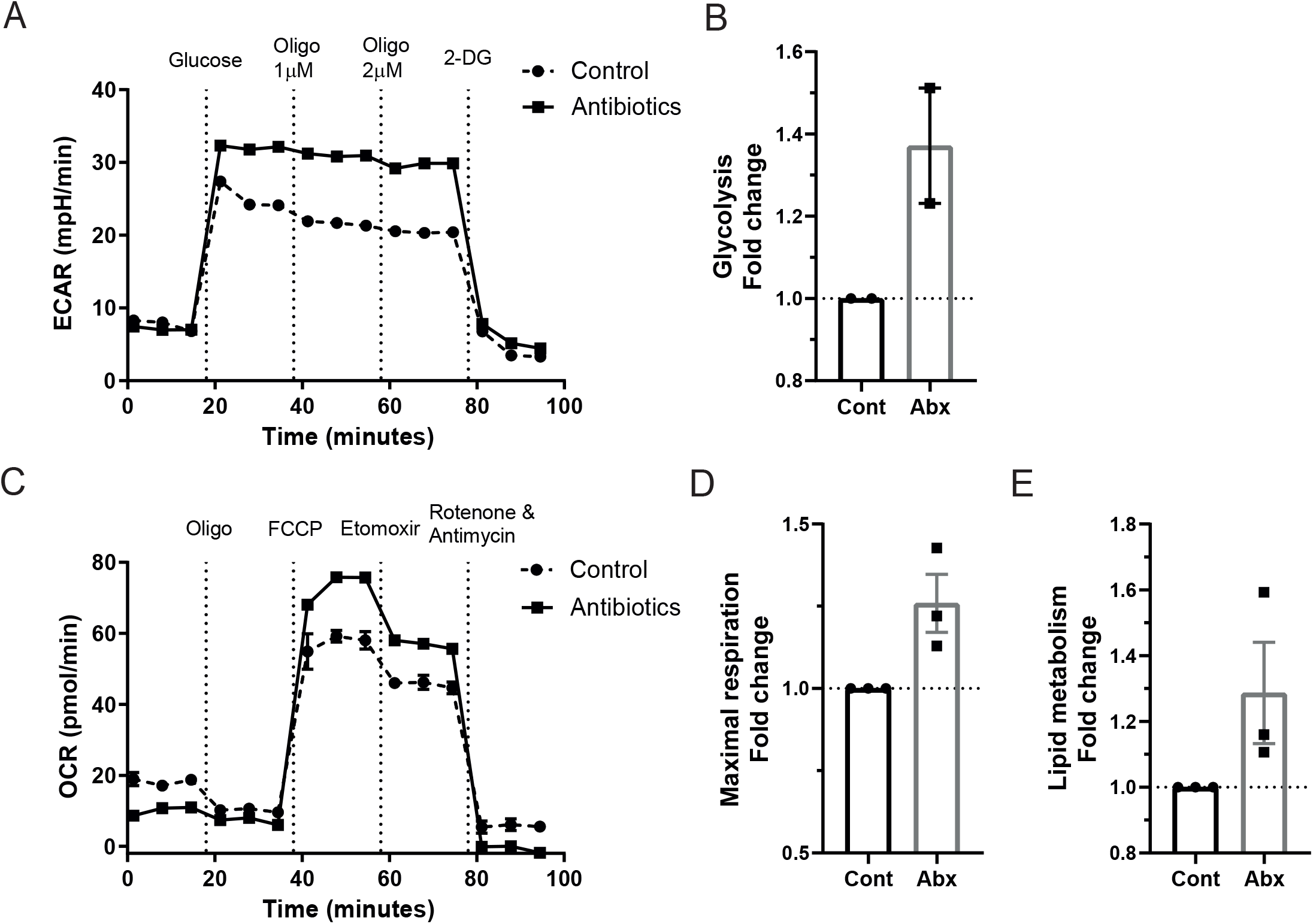
Seahorse analysis reveals increased metabolic output of large intestinal macrophages from antibiotic-treated mice. LiveCD45^+^SiglecF^−^Ly6G^−^CD11b^+^CD64^+^ large intestinal cells were sorted using flow cytometry from antibiotic and control treated mice and analysed by Seahorse technology. (**A**) Extracellular acidification rate (ECAR) was measured after addition of 25mM glucose, 1μM and 2μM oligomycin and 50mM 2-Deoxy-D-glucose (2-DG). Shown is a representative plot of two independent experiments. (**B**) ECAR fold change after addition of glucose (an indication of glycolysis). Fold change is relative to control samples. Average readings of 10 samples for each group from two independent experiments. (**C**) Oxygen consumption rate (OCR) was measured after addition of 2μM oligomycin, 1.5μM carbonyl cyanide-4-(trifluoromethoxy) phenylhydrazone (FCCP), 200μM etomoxir and 100nM rotenone and 1μM antimycin A. Data is representative of three independent experiments. (**D**) OCR fold change after addition of FCCP (an indication of maximal mitochondrial respiration). Fold change is relative to control samples. (**E**) OCR fold change after addition of etomoxir (an indication of lipid metabolism). Fold change is relative to control samples.

### Macrophages from the small intestine are less metabolically active than their colonic counterparts with reduced effects of the intestinal microbiota

The gastrointestinal tract varies considerably in function, environment, and composition of the local microbiota, with a reduced microbial load and diversity in the small intestine^29,30^. We assessed whether macrophages from the small intestine versus colon differ in their metabolic response to antibiotic treatment.

Small intestinal macrophages expressed lower levels of the amino acid transporter LAT1, with reduced CD98 protein (Fig. 4A) and Slc7a5 mRNA (Fig. 4B) compared to their colonic counterparts. Uptake of kynurenine (Fig. 4C) and the florescent glucose mimic 2-NBDG (Fig. 4D) was also reduced on macrophages in the small intestine, indicating restricted amino acid and glucose transport, respectively. Furthermore, small bowel macrophages had a lower mitochondrial mass (Fig. 4E), mitochondrial membrane potential (Fig. 4F) and uptake of fatty acid (Fig. 4G). Accordingly, antibiotic treatment had limited effects on metabolic functions of macrophages from the small intestine, with amino acid uptake, expression of CD98 and glucose uptake remained unchanged in contrast to macrophages from the colon (Fig. 4A, 4C-D). Finally, glycolysis and OXPHOS from CD11^+^CD64^+^ macrophages were analysed by Seahorse^TM^. ECAR of small intestinal macrophages was lower compared to colonic macrophages and indicated lower levels of glycolysis (Fig. 4H), while OCR was unchanged (Fig. 4I). Collectively, these data indicate that macrophages in the small intestine are less metabolically active than colonic macrophages and demonstrate reduced amino acid uptake, glycolysis, and lipid metabolism. Furtheremore, microbial-induced disruption of the microbiota had limited effects on macrophages in the small intestine.

**Figure 4:**
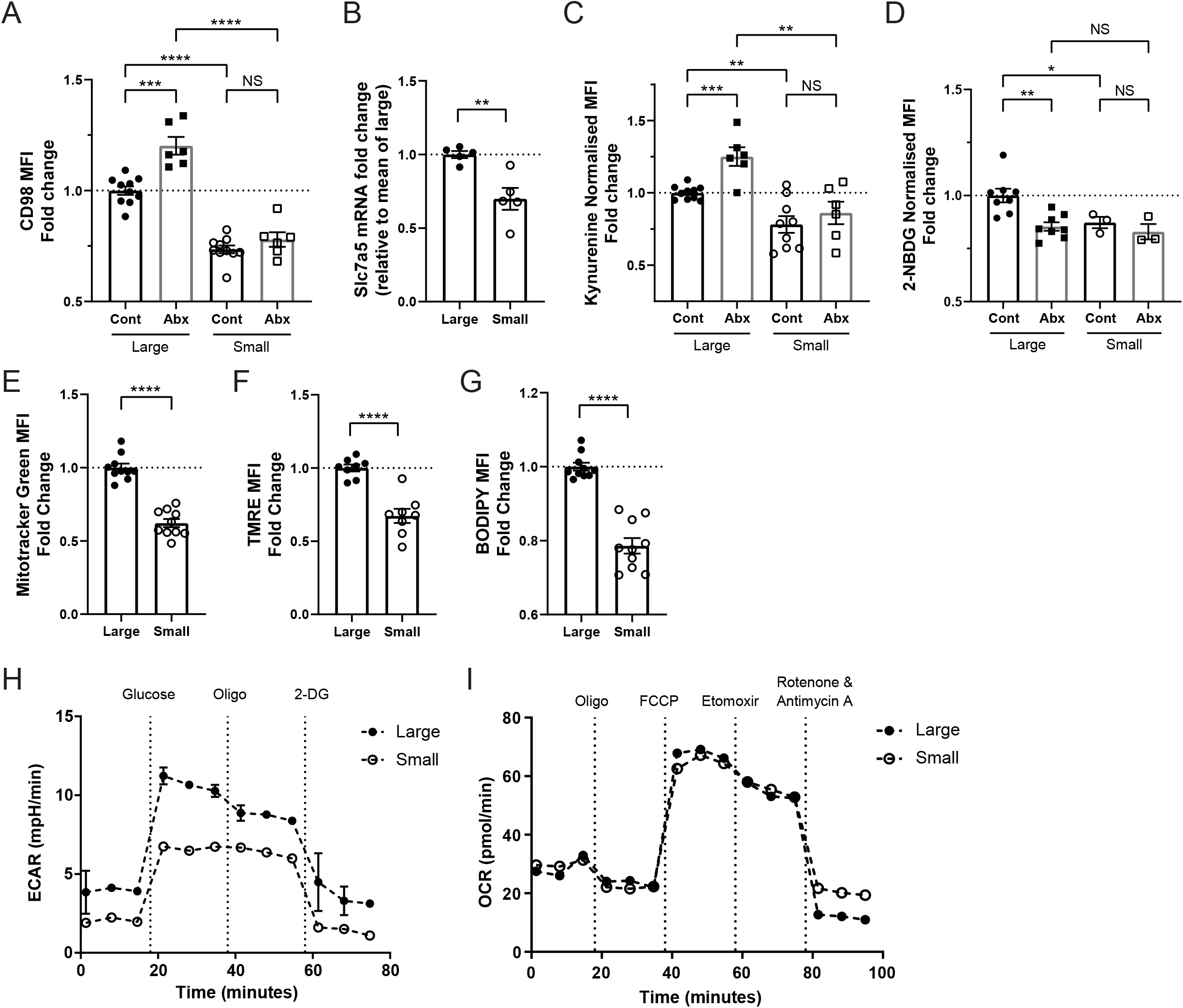
Small intestinal macrophages are less metabolically active, and less metabolically dependent on the gut microbiota. Mice were treated with antibiotic or control water and intestinal macrophages (Live CD45^+^Ly6G^−^CD11b^+^CD64^+^) were identified from the large or small bowel by flow cytometry. **(A)** Mean fluorescent intensity of CD98 protein expression on intestinal macrophages (Ly6C^−^MHC cII^+^). Fold change is normalised to mean of large control; n=6-10. **(B)** Relative mRNA expression of Slc7a5 on sorted Ly6C^−^ MHC cII^+^ macrophages. n=5. **(C)** Intestinal cell suspensions were treated with kynurenine and cell uptake measured on Ly6C^−^MHC cII^+^ intestinal macrophages at 450nm. MFI readings were normalised to background. Fold change is relative to the mean of the large control group; n=6-10. **(D-G)** Intestinal cell suspensions were treated with (**D**) 2-NBDG, (**E**) Mitotracker Green, (**F**) TMRE or (**G**) BODIPY FL C16 and cell uptake measured on Ly6C^−^ MHC cII^+^ intestinal macrophages at 530nm or 561nm. MFI readings were normalised to background. Fold change is relative to the mean of the large control groups; n=3-10. **(H)** Extracellular acidification rate (ECAR) was measured after addition of 25mM glucose, 2μM oligomycin and 50mM 2-Deoxy-D-glucose (2-DG). **(I)** Oxygen consumption rate (OCR) was measured after addition of 2μM oligomycin, 1.5μM carbonyl cyanide-4-(trifluoromethoxy) phenylhydrazone (FCCP), 200μM etomoxir and 100nM rotenone and 1μM antimycin A. *p<0.05, **p<0.01, ***p<0.001, ****p<0.0001, NS: non-significant, Student’s t-test.

### Lipid uptake defines intestinal tissue-resident macrophages

Given the limited effects of antibiotic-induced microbial disruption on tissue resident subsets of intestinal macrophages with slow turnover from the circulation in terms of induction of pro-inflammatory properties^19^, we aimed to investigate whether subsets of intestinal macrophages exhibited differences in their metabolic capacities in response to antibiotics. Macrophage subsets were characterised as replenished (CD4^−^Tim4^−^ and CD4^+^Tim4^−^ macrophages) or tissue-resident (CD4^+^Tim4^+^) in the colon. Although mitochondrial mass (Fig. 5A) and membrane potential (Fig. 5B) were comparable between the CD4^−^Tim4^−^, CD4^+^Tim4^−^, CD4^+^Tim4^+^ macrophages within the small intestine and colon, uptake of lipids as measured by BODIPY C16 was significantly higher in CD4 Tim4 double positive macrophages with the slowest turnover rate (Fig. 5C), in both compartments. To confirm these findings, mRNA was isolated from sorted Tim4 CD4 macrophage subsets and lipid metabolism genes quantified. Accordingly, expression of fatty acid uptake genes including the scavenger receptor *Cd36* and the primary macrophage fatty acid transporter *Slc27a1* increased in the CD4^+^Tim4^+^ resident macrophages (Fig. 5D). In contrast to this, *Fasn*, the primary fatty acid synthase, was downregulated in the double positive population (Fig. 5D). These data suggest lipid/fatty acid uptake (and not de nova synthesis) may be a requirement for gut macrophages for long-lived tissue residency, a characteristic of other tissue resident immune cells^31^, and likely have implications for macrophage function.

**Figure 5:**
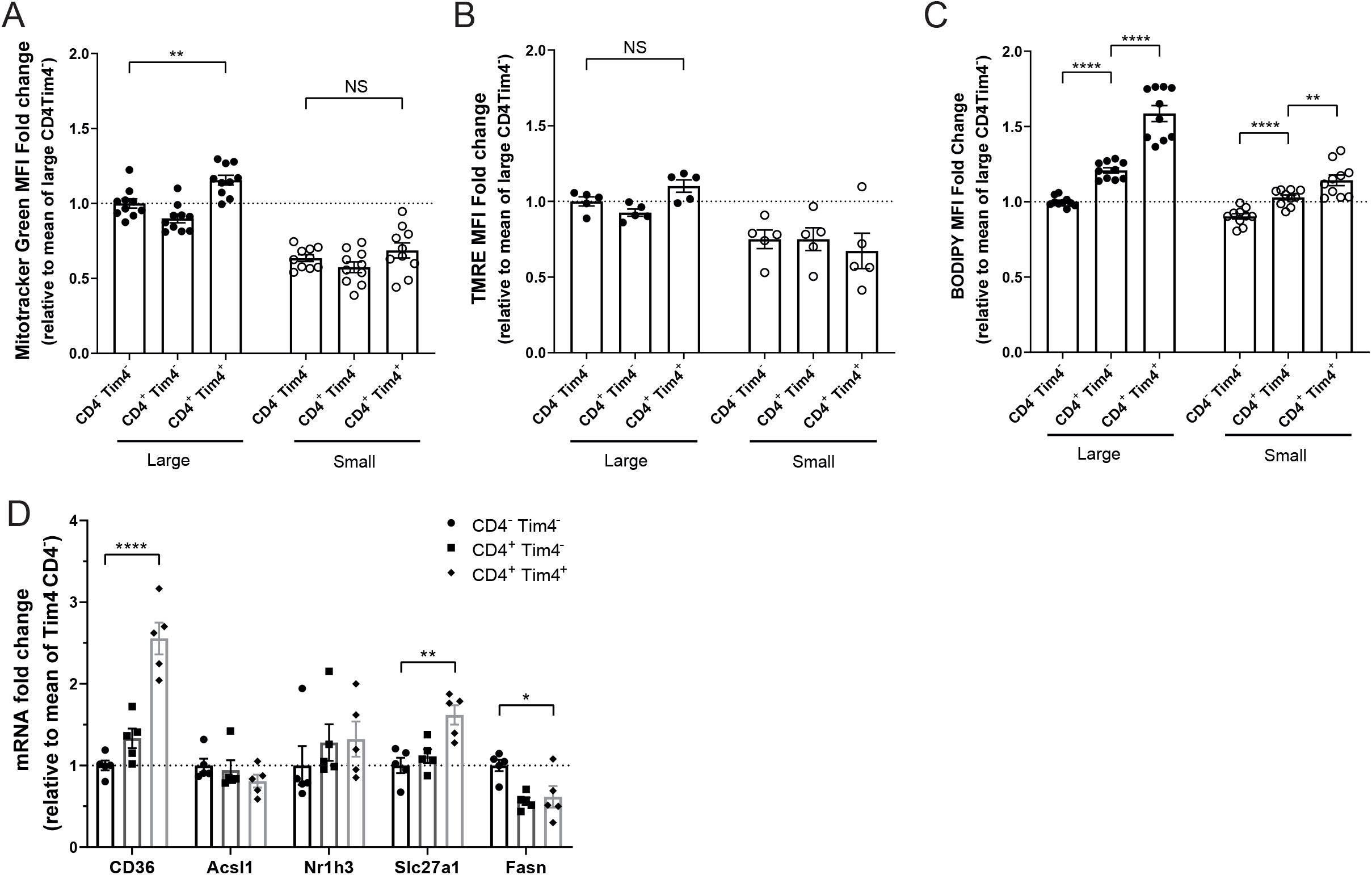
Fatty acid uptake defines the metabolism of intestinal tissue resident macrophages. Large and small bowel cell suspensions from naïve mice were isolated and Tim4^−^ CD4^−^, Tim4^−^ CD4^+^ and Tim4^+^ CD4^+^ macrophages identified by flow cytometry. **(A)** Cells were treated with 50nM Mitotracker Green and staining measured at 530nM. n=10. **(B)** Cells were treated with 25nM of TMRE and fluorescence measured at 586nM. n=5. **(C)** Cells were treated with 5μM BODIPY FL C16 in fatty acid free media, and uptake measured at 530nM, n=10. **(D)** Large intestinal macrophages (LiveCD45^+^SiglecF^−^Ly6G^−^CD11b^+^CD64^+^) were sorted by flow cytometry into Tim4^−^ CD4^−^, Tim4^−^ CD4^+^ and Tim4^+^ CD4^+^ subsets and mRNA expression of lipid uptake and synthesis-related genes was measured. n=5. *p<0.05, **p<0.01, ***p<0.001, ****p<0.0001, NS: non-significant, Student’s t-test.

## DISCUSSION

We demonstrate that intestinal macrophages can constitutively take up amino acids from their local environment and that this is regulated by the intestinal microbiota. Antibiotic-induced microbial disruption further enhanced amino acid uptake and as such, increased the metabolic capacities of macrophages in the intestine. Taken with the direct association between cellular metabolism and activation^6,24,27,32,33^, these data provide novel insight into pathways by which macrophage activation in the intestine may be regulated by the commensal microbiota to avoid inappropriate inflammation. The reduced metabolic capacity of macrophages in the small intestine is likely to be due to the environmental differences between the different regions of the intestine^29,30^ but has important implications for macrophage function. The small bowel is responsible for the absorption and digestions of protein, monosaccharides and vitamins; hence immune cell homeostasis is more dependent on diet in the small intestine compared to the colon^34^. Yet the restricted metabolic capacity of small intestinal macrophages compared to their colonic counterparts indicate that the intestinal microbiota may play a bigger role in shaping macrophage metabolism in the intestine than diet – despite the obvious implications for availability of amino acids from dietary protein in the small intestine.

We have previously demonstrated that intestinal macrophages from antibiotic treated mice exhibit pro-inflammatory properties, responding more potently to bacterial-derived LPS and expressing enhanced levels of iNOS, aligning with classically activated macrophages^19^. Although classically activated macrophages rely on glycolysis for energy production to enable rapid responses to pathogens when needed^35^, we show here that antibiotic-induced microbial disruption increases both glycolysis and OXPHOS pathways. Traditionally, a characteristic of alternatively activated macrophages is their utilisation of OXPHOS for efficient energy production where downstream products of the TCA cycle such as itaconate inhibit inflammation through suppression of IL-6 and IL-1b^36,37^. However, despite upregulation of both metabolic pathways, antibiotic treatment induced lower levels of alternative activation marker CD206 and higher amounts of CD11c a classical activation marker whilst increasing macrophage activation as indicated by phagocytic activity, again supporting induction of a pro-inflammatory phenotype in macrophages by antibiotics as seen in our previous studies.

Our data support a strong role for the microbiota in limiting macrophage metabolism in the intestine, in contrast to regulation of macrophage metabolism at other mucosal sites e.g. the lung where the microbiota plays a limited/absent role but cellular metabolism is thought to be shaped by local glucose levels^38^. However, those data also indicate aberrant control of pulmonary macrophage metabolism can result in upregulation of both glycolysis and OXPHOS pathways.

Our observation that Tim4^+^ CD4^+^ tissue resident intestinal macrophages upregulate fatty acid transporters and scavengers whilst downregulating fatty acid synthesis genes indicate that this macrophage subset may utilise uptake of local fatty acids for function. Indeed, CD36 and has been defined as a key part of metabolic reprograming that occurs when CD8 memory T cells become resident in the skin^31,39^, suggesting CD36 may be involved in tissue residency. Uptake of lipids and their metabolism through beta-oxidation in mitochondria grants these cells longevity in skin tissue. These differences in metabolic function between macrophage subsets in the intestine are likely to impact on their function; as yet it is unclear what the functional differences between these recruited versus resident macrophages in the intestine are. Given CD36 expression is correlated with an immunosuppressive phenotype seen in tumour-associated macrophages (TAM)^40^, taken with fatty acid uptake being a hallmark feature of alternatively activated macrophages^41^, our data suggest that the functions of CD4^+^Tim4^+^ tissue resident macrophages may be more directed towards tissue remodelling and integrity rather than response to pathogens. Indeed, Tim4^+^CD4^+^ macrophages produce lower levels of pro-inflammatory type 1 cytokine IL-6 compared to their Tim4^−^CD4^−^ recruited counterparts^4^. Further work is needed to elucidate the roles of specific macrophage subsets in intestinal immune homeostasis.

## METHODS

### Mice

WT C57BL/6 mice (Charles River) were maintained under specific pathogen-free conditions at the University of Manchester, UK. Age- and gender-matched adult (8-12 week old, female) animals were used in all experiments and protocols approved by the University of Manchester Animal Welfare Ethical Review Boards. All experiments were carried out under license issued by and according to U.K. Home Office regulations.

### Antibiotic treatment

Mice were treated with an antibiotic cocktail containing ampicillin (1g/L), metronidazole (1g/L), neomycin (1g/L), gentamicin (1g/L) and vancomycin (0.5g/L) in drinking water for seven days, which was replenished once on day three or four.

### Isolation of intestinal lamina propria cells

Large and small intestines were removed from mice, cleaned with PBS and chopped into 5mm sections before incubation with 2mM EDTA in HBSS three times for 15 minutes on shake at 210rpm. After washing, tissue was digested with 1.25mg/mL collagenase D (Sigma), 0.85mg/mL collagenase V (Sigma) and 1mg/mL dispase (Gibco, Invitrogen) in 10% FBS RPMI-1640 (Sigma) at 37°C for 30 mins (small intestine) or 35 mins (large intestine).

### Faecal metabolite analysis

Performed as described previously^19^. Briefly, stools were prepared for ^1^H nuclear magnetic resonance (NMR) spectroscopy and recorded at 600MHz on a Bruker Avance spectrometer (Bruker BioSpin GmbH, Rheinstetten, Germany) running Topsin 2.0 software and fitted with a broadband inverse probe. All metabolites were quantified using Chenomx NMR suite 7.6 software and by use of the 2D-NMR methods, COSY, HSQC and HMBC.

### Flow cytometry and sorting of cells

Isolated cells were stained with Fc block (BD Sciences) for 5 minutes prior to staining at 4°C in the dark using the antibodies listed in supplementary table 1 and analysed using an LSR Fortessa cytometer (BD Biosciences) and FlowJo software (BD). Monocytes and macrophages were sorted using a FACS Aria Fusion, as live gated CD45^+^SiglecF^−^Ly6G^−^ CD11b^+^CD64^+^ cells to >97% purity and further subdivided into subsets based on MHC class II, Ly6C, Tim4 and CD4 expression.

### SCFA and amino acid treatment

Mice were administered acetate (67.5mM), propionate (25.9mM) and butyrate (40mM) in drinking water (concentrations reflective of ratios within the intestine^42^) with or without antibiotics for seven days. For the amino acid supplementation experiments, mice received 100mM each of lysine, leucine and glutamine in drinking water with or without antibiotics for seven days.

### Amino acid uptake assay

Based on the kynurenine uptake assay devised by Linda Sinclair and Doreen Cantrell^21^. After staining to distinguish macrophages, each sample was split into four tubes: HBSS alone (FMO/background), L-kynurenine (Sigma, 200uM final), L-kynurenine + leucine (Sigma, 5mM final) and kynurenine + 2-amino-2-norbornanecarboxylic acid (BCH) a LAT1 inhibitor, (Sigma, 10mM final). Samples were placed at 37°C for 5 minutes, before addition of Cytofix Fixation Buffer (BD Biosciences), pulse vortex and incubation at 4°C for 20 minutes. After washing, cells were analysed by flow cytometry. Kynurenine was detected on the e450/BV421 channel (BV450/50 filter on LSR Fortessa).

### Phosphoflow

Cell suspensions were lysed, fixed and stained using a modified BD Phosphoflow protocol. After addition of prewarmed lyse/fix buffer (BD Phosphoflow) cells were inverted 10 times and incubated at 37°C for 10 minutes. Cells were washed and centrifuged 600g for six minutes twice before permeabilising with Perm Buffer III for 30 minutes at 4°C. After washing cells were treated with Fc block for 5 mins before staining with phosphoflow and regular antibodies (see antibody list, supplementary table 1).

### Cellular metabolism assays

Intestinal cell suspensions were stained with an antibody cocktail to distinguish macrophages before addition of Mitotracker Green (50nM final) and Tetramethylrhodamine, ethyl ester (TMRE) (25nM final) in 10% FBS RPMI-1640 and incubation for 30 mins at 37 °C with CO_2_. Cells were washed twice before analysis by flow cytometry. Mitotracker Green was detected on the FITC channel (B530/30 filter on LSR Fortessa) and TMRE on the PE channel (Y568/15 filter on LSR Fortessa). For fatty acid uptake, 5uM BODIPY FL C16 (Invitrogen) in 0.5% fatty acid-free BSA RPMI-1640 (both Sigma) was added to cells for 10 minutes in a 37 °C 5% CO_2_ incubator and signal detected on the FITC channel (B530/30 filter on the LSR Fortessa). For glucose uptake, 100uM 2-NBDG was added to cells in glucose-free RPMI 1640 media (Gibco, ThermoFisher Scientific) for one hour at 37°C in an incubator and detected on the FITC channel (B530/30 filteron the LSR Fortessa).

### Quantitation of gene expression by real time PCR

Total RNA was extracted from monocytes and macrophages from the intestines of individual or pooled samples using the RNeasy Micro Kit (Qiagen). RNA were reverse transcribed to cDNA using the High Capacity RNA-to-cDNA Kit (Applied Biosystems, Thermo Scientific) and gene expression was assayed using quantitative reverse transcription PCR using PerfeCTa SYBR Green Fastmix Low ROX (Quanta) on the QuantStudio 12K Flex (Applied Biosystems) with primers (Integrated DNA Technologies) listed in supplementary table 2.

### Phagocytosis assay

After digestion, intestinal cell suspensions were placed in 10% FBS RPMI-1640 in a 5% CO_2_ 37°C incubator to allow time for attachment to the tubes for 1.5-2 hours. After washing, 100uL of 1mg/mLpHrodo Red E.coli BioParticles (Invitrogen) were added in PBS and cells placed at 4°C (negative control) or 37°C for 30 minutes. After washing, cells were stained with the macrophage antibody panel and analysed by flow cytometry. pHrodo Red BioParticles were detected on the PE channel (Y568/15 filter, LSR Fortessa).

### Seahorse

Extracellular acidification rates (ECAR) and oxygen consumption rates (OCR) were measured using an XF-96 Extracellular Flux Analyser (Seahorse Bioscience, Agilent Tecnologies). ECAR was measured at baseline and after addition of 25mM glucose, 1-2uM oligomycin and 50mM 2-deoxy-d-glucose (2-DG). OCR were taken under basal conditions and following the addition of 1-2uM oligomycin, 1.5uM fluro-carbonyl cyanide phenylhydrazone (FCCP), 200uM etomoxir and 100nM rotenone +1uM antimycin A (all purchased from Sigma).

## ACKNOWLEDGEMENTS

The authors thank M. Jackson, D. Chapman and G.Howell of the FBMH flow cytometry facility for cell sorting, and members of the BSF for animal husbandry, in particular R. Hallworth, R. Hodgkiss, M. Jackson and N. Locke. This research was funded by the Wellcome Trust [206206/Z/17/Z (to E.R.M.)]

## AUTHOR CONTRIBUTIONS

NAS and ERM wrote the manuscript and designed the study. NAS, MAL, LJH, ERM, RJH performed experiments and analysed data.

## SUPPLEMENTRY TABLES

**Table S1.**
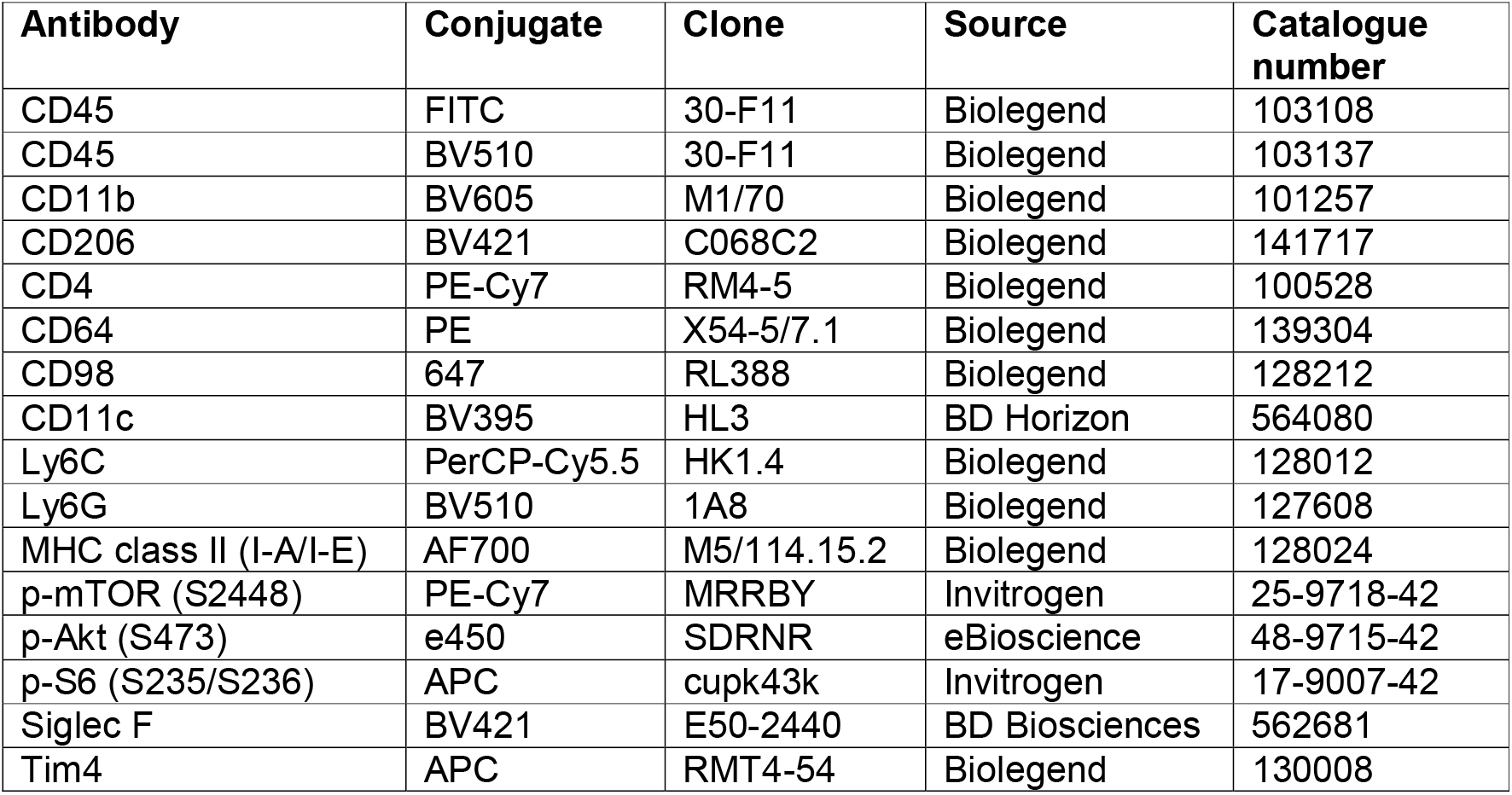
Antibodies.

**Table S2.**
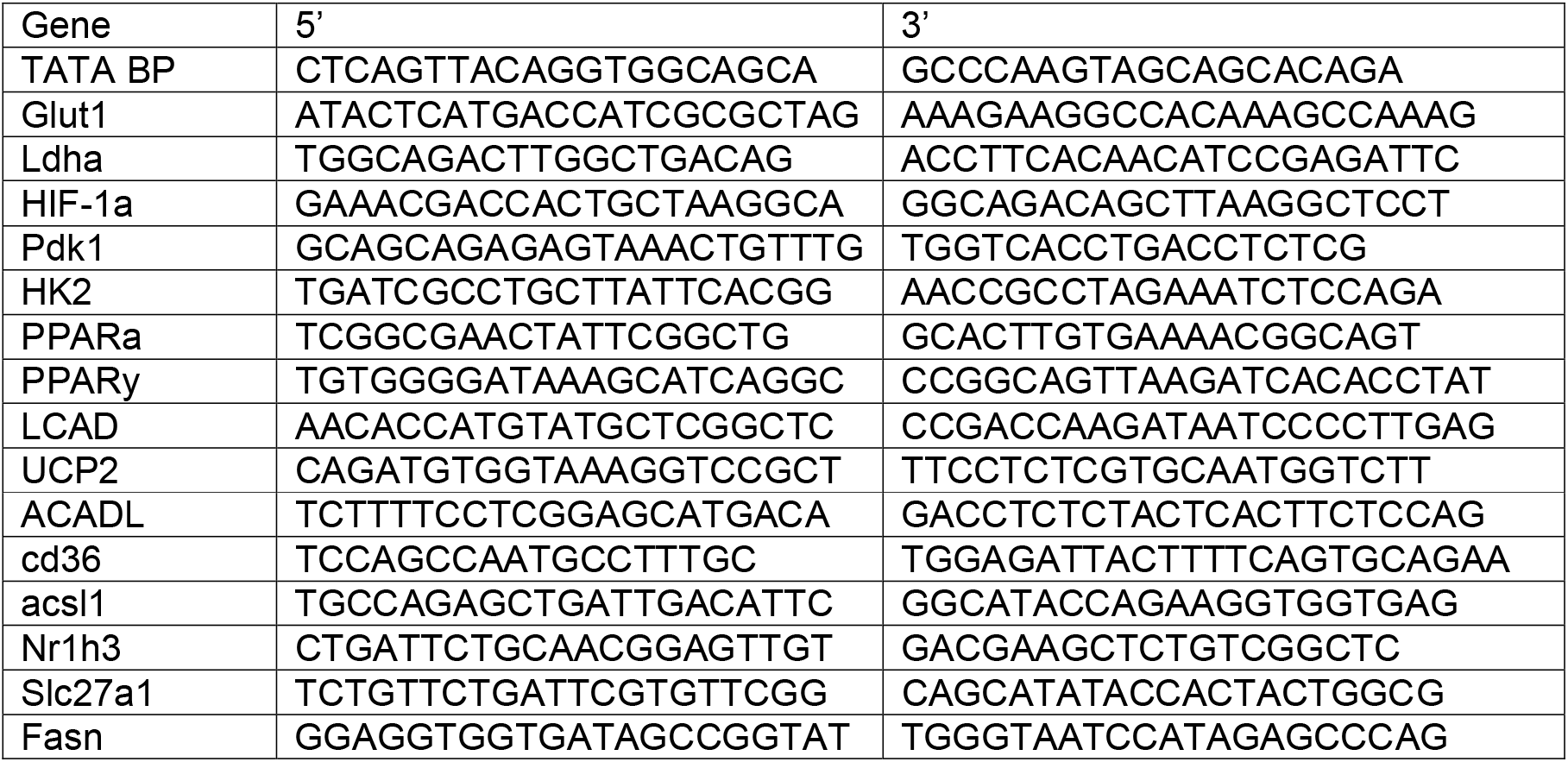
Primers sequences for gene expression.

